# Post-embryonic hourglass patterns mark ontogenetic transitions in plant development

**DOI:** 10.1101/035527

**Authors:** Hajk-Georg Drost, Julia Bellstäedt, Diarmuid S. Ó’Maoiléidigh, Anderson T. Silva, Alexander Gabel, Claus Weinholdt, Patrick T. Ryan, Bas J.W. Dekkers, Leónie Bentsink, Henk Hilhorst, Wilco Ligterink, Frank Wellmer, Ivo Grosse, Marcel Quint

**Author notes:** present address: Max Planck Institute for Plant Breeding Research, 50829 Cologne, Germany (Diarmuid S. Ó’Maoiléidigh).

## Abstract

The historic developmental hourglass concept depicts the convergence of animal embryos to a common form during the phylotypic period. Recently, it has been shown that a transcriptomic hourglass is associated with this morphological pattern, consistent with the idea of underlying selective constraints due to intense molecular interactions during body plan establishment. Although plants do not exhibit a morphological hourglass during embryogenesis, a transcriptomic hourglass has nevertheless been identified in the model plant *Arabidopsis thaliana*. Here, we investigated whether plant hourglass patterns are also found post-embryonically. We found that the two main phase changes during the life cycle of *Arabidopsis*, from embryonic to vegetative and from vegetative to reproductive development, are associated with transcriptomic hourglass patterns. In contrast, flower development, a process dominated by organ formation, is not. This suggests that plant hourglass patterns are decoupled from organogenesis and body plan establishment. Instead, they may reflect general transitions through organizational checkpoints.

## Introduction

Based on von Baer’s third law of embryology (von Baer 1828), it has been observed that mid-stage embryos of animal species from the same phylum share morphological similarities. Because these embryos tend to be more divergent at early and late embryogenesis, this morphological pattern has been termed the ‘developmental hourglass’ (Duboule 1994; Raff 1996) (Fig. 1a). The window of maximum morphological conservation in mid-embryogenesis coincides with the onset of organogenesis during body plan establishment and is called phylotypic stage (Sander 1983) or phylotypic period (Richardson 1995, Kalinka et al. 2010). It has been suggested that a likely cause for this conservation is a web of complex interactions among developmental modules (e.g. organ primordia) during body plan establishment, which results in selective constraints that minimize morphological divergence (Raff 1996) (Fig. 1a). While controversially debated for decades, in recent years the concept of the developmental hourglass has been largely confirmed at the transcriptomic level. Several studies showed that the degree of sequence conservation, the phylogenetic age of transcriptomes, or the similarity of gene expression profiles maximize during the phylotypic period (Hazkani-Covo et al. 2005; Irie and Sehara-Fujusawa 2007; Artieri et al. 2009; Cruickshank and Wade 2008; Kalinka et al. 2010; Domazet-Lošo and Tautz 2010; Irie and Kuratani 2011; Levin et al. 2012; Wang et al. 2013), which is in agreement with a potentially causative association with body plan establishment.

**Figure 1.**
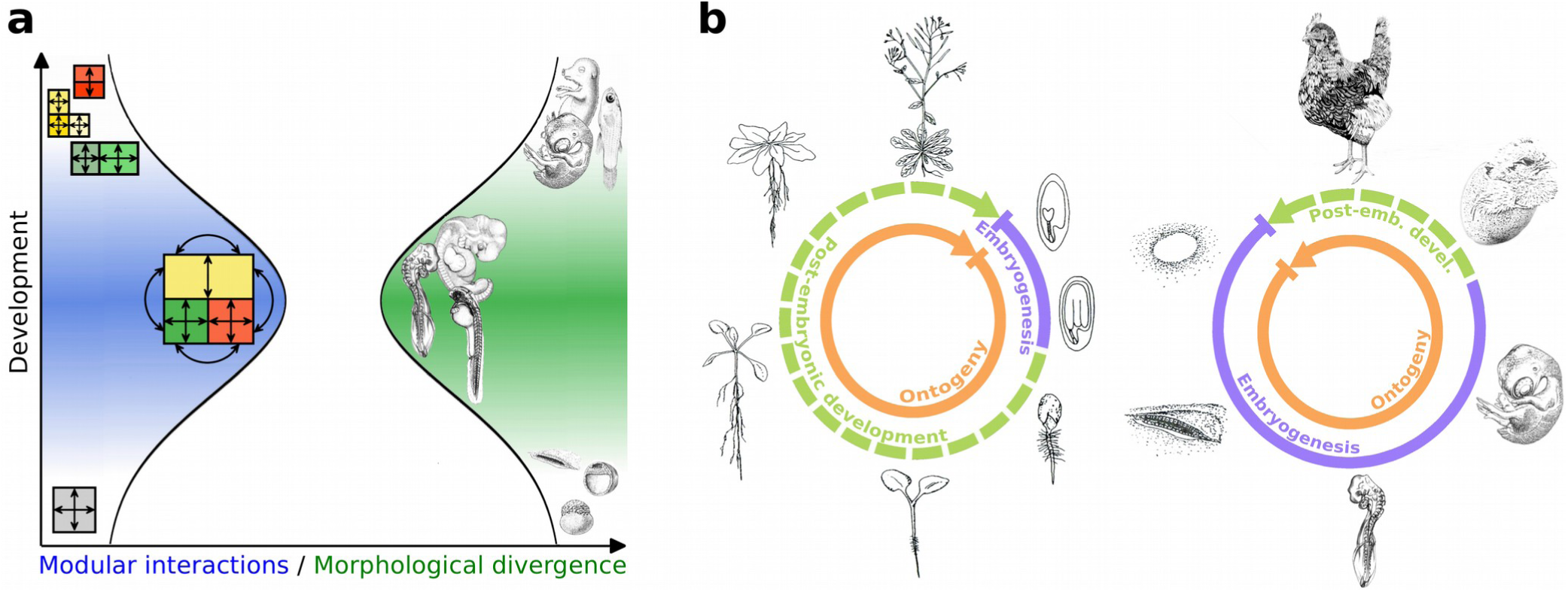
The developmental hourglass model in the context of differences in plant and animal development. (**A**) According to Raff (Raff 1996), a web of complex interactions among developmental modules results in selective constraints during mid embryogenesis. In the phylotypic period modular interactions maximize and morphological divergence minimizes resulting in the bottleneck of the developmental hourglass model (illustration adapted from (Irie and Kuratani 2011)). (**B**) The part of the ontogenetic life cycle that is covered by embryogenesis varies dramatically between plants and animals. Mature plant embryos have a limited number of organs and little complexity. Most organs develop post-embryonically. In contrast to animals, the plant body plan is not fixed. It constantly changes in response to the environment. Animal development is largely embryonic. Mature animal embryos often reach a level of complexity that is comparable to adult individuals.

In contrast to animals with their almost exclusively embryogenic development, organ formation in plants occurs largely post-embryonically (Fig. 1b). Hence, a web of comparably complex modular interactions between developing organ primordia, which might underly the selective constraints during the phylotypic period in animals, is possibly never achieved during plant embryogenesis. However, a transcriptomic hourglass pattern has nonetheless been observed for plant embryogenesis (Quint et al. 2012; Drost et al. 2015a) (as well as for fungal development (Cheng et al. 2015)), indicating that it may not be causally connected to organogenesis, as suggested by the animal model. We therefore wondered whether in plants these patterns might instead be associated with developmental transitions. Embryogenesis can be viewed as such a transition, namely from a single-celled zygote to a complex, multi-cellular embryo. To test this hypothesis, we generated transcriptomic data sets that cover the two most important ontogenetic transitions in post-embryonic development in *A. thaliana:* the transition from the embryonic to the vegetative phase, and the transition from the vegetative to the reproductive phase. As a control, we also analyzed a transcriptomic time series for flower development, a process that is dominated by organogenesis. We then performed phylotranscriptomic analyses (Domazet-Lošo and Tautz 2010; Quint et al. 2012; Drost et al. 2015a; Drost et al. 2015b), which assess the phylogenetic age of transcriptomes expressed over sequential developmental stages (Supplementary Figure S1), and tested the resulting profiles for the characteristic hourglass shape. If indeed, post-embryonic developmental processes would be governed by hourglass patterns, this would suggest that hourglass patterns are not restricted to embryogenesis and possibly a wide-spread phenomenon that governs multiple processes. Furthermore, the potentially causative relationship between organogenesis, body plan establishment and hourglass patterns would need to be reevaluated.

## Results and Discussion

To study the transition from embryogenesis to the vegetative phase, we generated transcriptomic information for seven sequential ontogenetic stages during seed germination. The stages sampled included mature dry seeds, six-hours imbibed seeds, seeds at testa rupture, radicle protrusion, root hair (collet hair) appearance, the appearance of greening cotyledons, and established seedlings with fully opened cotyledons (Fig. 2a; Supplementary Fig. S2). We then combined the transcriptomic information with previously generated gene age information (Drost et al. 2015). Based on an age-assignment approach called phylostratigraphy (Domazet-Lošo et al. 2007) (Supplementary Fig. S1), genes can be sorted into discrete age categories named phylostrata (PS) (Domazet-Lošo et al. 2007). For *A. thaliana*, we defined 12 age classes ranging from old (PS1) to young (PS12). Next, we computed the transcriptome age index (TAI) (Domazet-Lošo and Tautz 2010) for each developmental stage, which is defined as the weighted mean of gene ages using the stage-specific expression levels as weights. The TAI therefore describes the phylogenetic age of a transcriptome.

**Figure 2.**
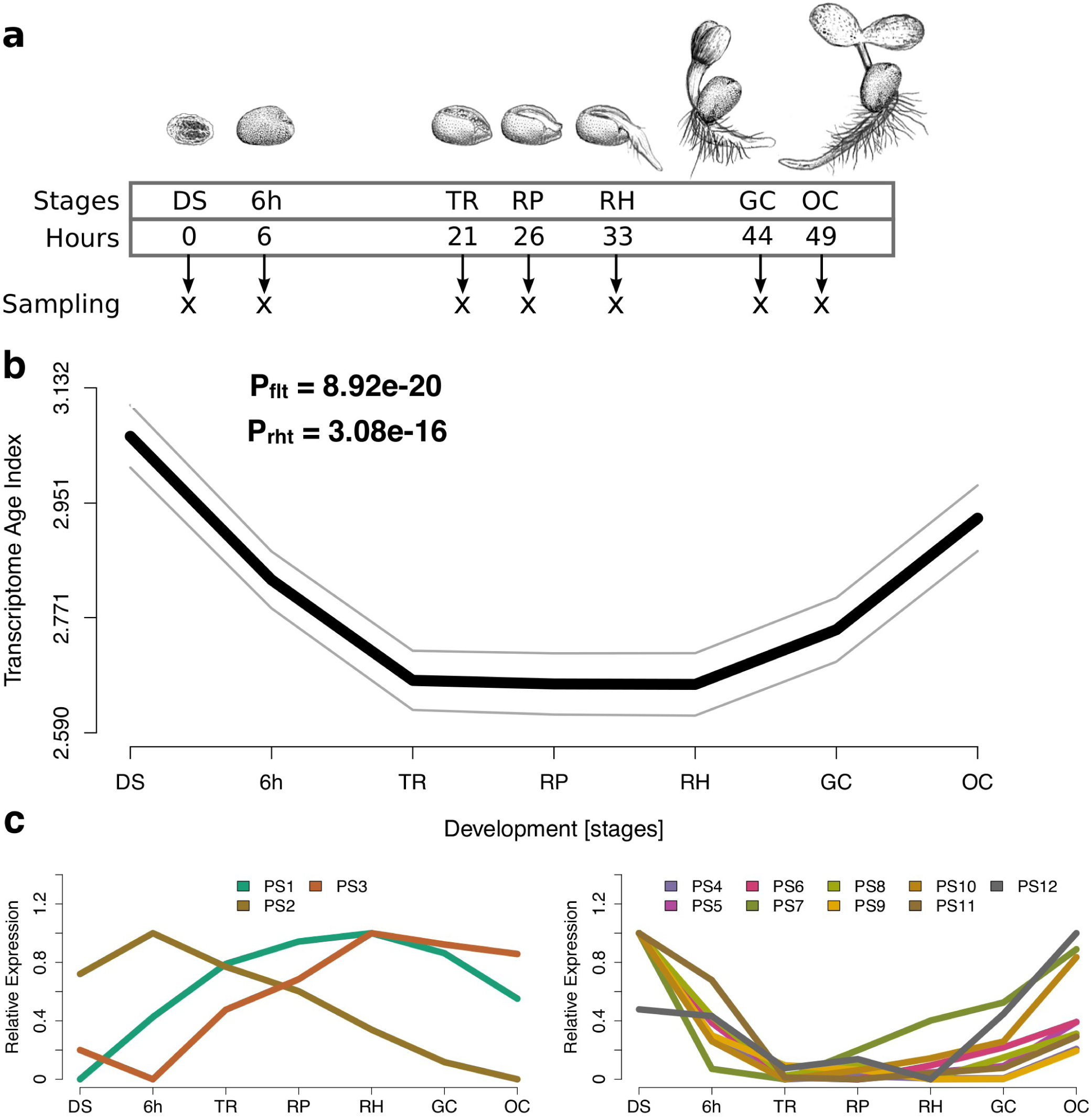
Transcriptome age index analysis for germination in *A. thaliana*. (**A**) Illustration of the developmental stages for which transcriptome data were generated. (**B**) The transcriptome age index (TAI) profile across germination follows an hourglass-like pattern. The gray lines represent the standard deviation estimated by permutation analysis. P-values were derived by application of the flat line test (Drost et al. 2015a) (P_flt_) and the reductive hourglass test (Drost et al. 2015a) (P_rht_). (**C**) Relative expression levels for each phylostratum (PS) separately. The stage with the highest mean expression levels of the genes within a PS was set to relative expression level = 1, the stage with the lowest mean expression levels of the genes within a PS was set to relative expression level = 0, the remaining stages were adjusted accordingly. PS were classified into two groups: Group ‘old’ contains PS that categorize genes that originated before complex/multicellular plants evolved (PS1-3) and Group ‘young’ contains PS that categorize genes that originated after complex plants evolved (PS4-12). (**A-C**) DS, mature dry seeds; 6h, six-hours imbibed seeds; TR, seeds at testa rupture; RP, radicle protrusion; RH, appearance of the first root hairs; GC, appearance of greening cotyledons; OC, fully opened cotyledons.

As shown in Figure 2b, the TAI profile for the embryonic-to-vegetative phase transition displays an hourglass pattern with high TAI values at early and late stages and low TAI values at intermediate stages. We confirmed this observation through statistical tests (flat line test (Drost et al. 2015a): P = 8.92e-20; reductive hourglass test (Drost et al. 2015a): P = 3.08e-16; Supplementary Fig. S3a). The waist of the hourglass corresponded to the phylogenetically oldest transcriptomes stemming from the ‘testa rupture’ to ‘radicle protrusion’ stages. These stages mark the emergence of the seedling from the seed, likely the transition period of this process, at which germination becomes irreversible (Fig. 2b). We finally also studied the relative expression levels of genes of different phylostrata and found that the hourglass pattern is caused by a largely antagonistic behavior of old and young genes (Fig. 2c), similar to what has been previously reported for embryogenesis (Quint et al. 2012; Drost et al. 2015a).

**Figure 3.**
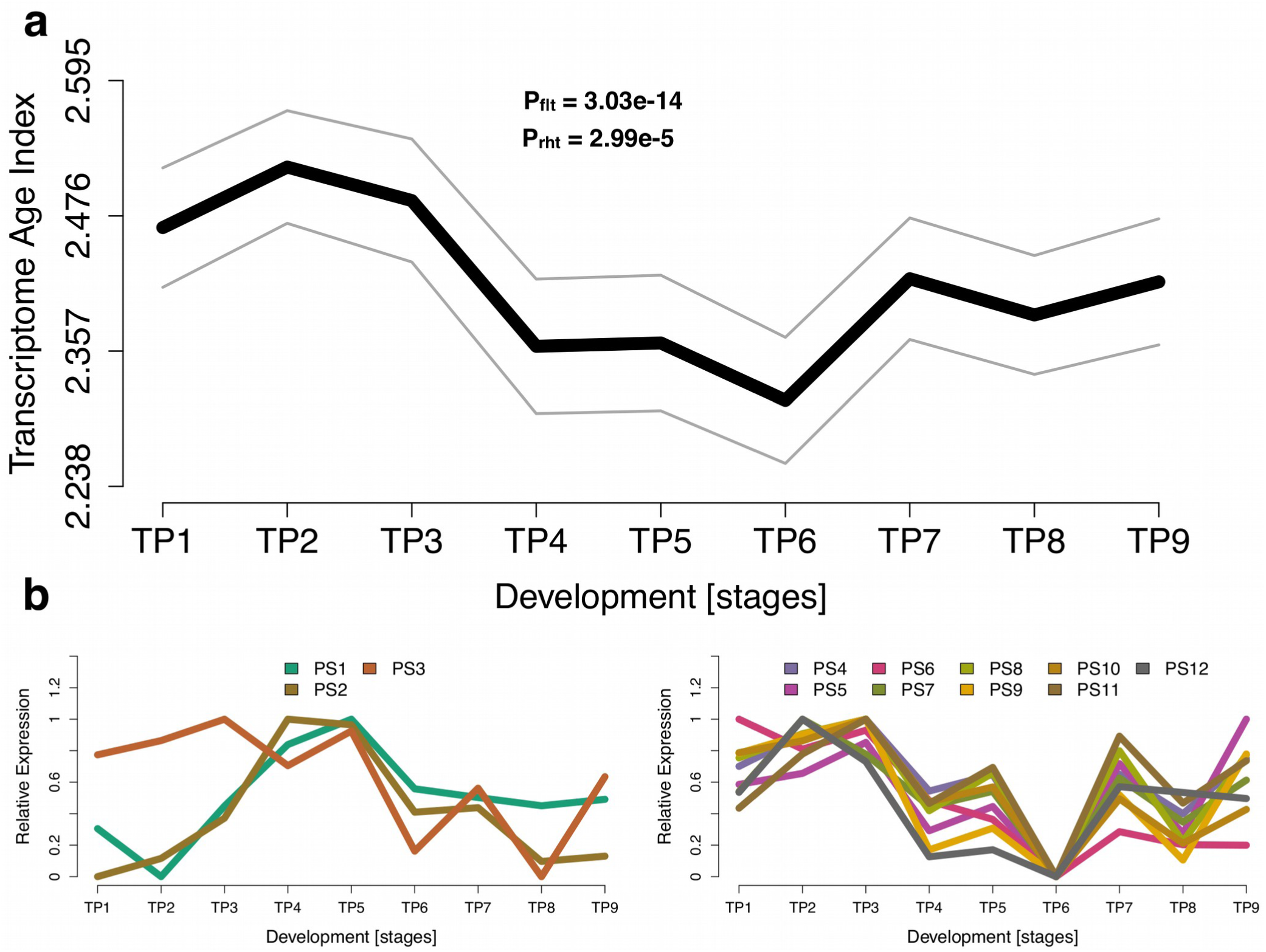
Transcriptome age index analysis for the transition from vegetative to reproductive growth in *A. thaliana*. (**A**) The transcriptome age index (TAI) profile across the transition to flowering follows an hourglass-like pattern. The gray lines represent the standard deviation estimated by permutation analysis. P-values were derived by application of the flat line test (Drost et al. 2015a) (P_flt_) and reductive hourglass test (Drost et al. 2015a) (P_rht_). (**B**) Relative expression levels for each phylostratum (PS) separately. The stage with the highest mean expression levels of the genes within a PS was set to relative expression level = 1, the stage with the lowest mean expression levels of the genes within a PS was set to relative expression level = 0, the remaining stages were adjusted accordingly. PS were classified into two groups: Group ‘old’ contains PS that categorize genes that originated before complex / multicellular plants evolved (PS1-3) and Group ‘young’ contains PS that categorize genes that originated after complex plants evolved (PS4-12). (**A-B**) TP: time point; TP1: 1 day after shift to long day photoperiods (LD), TP2: 2 days after shift to LD, TP3: 3 days after shift to LD, TP4: 4 days after shift to LD, TP5: 5 days after shift to LD, TP6: 6 days after shift to LD, TP7: 7 days after shift to LD, TP8: 8 days after shift to LD, TP9: 9 days after shift to LD.

We next tested whether a transcriptomic hourglass pattern also underlies the vegetative-to-reproductive phase transition. During this so-called floral transition, the leaf-producing shoot apical meristem is converted into an inflorescence meristem, which forms flowers (Huijser and Schmid 2011). Morphologically, completion of the floral transition can be observed by the bolting inflorescence. However, as the actual transition occurs several days before bolting, we also assessed the expression of floral homeotic genes and other marker genes to better map the time of transition to the reproductive state (Supplementary Fig. S4). Based on this information, we synchronized flowering time in the sampling population (Supplementary Fig. S5; see methods) and generated transcriptome data from the shoot apex before, during, and after floral transition.

Figure 3a shows the results from the TAI analysis for nine samples covering the floral transition. We identified a robust hourglass pattern (reductive hourglass test (Drost et al. 2015a): P = 2.99e-5, Fig. 3a, Supplementary Fig. S3b) that significantly deviated from a flat line (flat line test (Drost et al. 2015a): P = 3.03e-14). Similar to embryogenesis (Quint et al. 2012; Drost et al. 2015a) and seed germination (Fig. 2c), analysis of relative expression levels of genes assigned to different age classes revealed a largely antagonistic behavior of old and young genes (Fig. 3b).

Taken together, these observations demonstrate that in plants not only embryogenesis, but also the embryo-to-vegetative and vegetative-to-reproductive phase transitions progress through a stage of evolutionary conservation with older transcriptomes being active in mid development. Thus the hourglass pattern, which was previously discussed only with regard to embryogenesis, appears to be more widespread, at least in plants. In fact, the embryonic hourglass is possibly only one of many developmental processes governed by hourglass patterns.

Because no new organs are established during the two post-embryonic phase transitions assessed here, our results also support the aforementioned conjecture that transcriptomic hourglass patterns are not specifically associated with organogenic processes. To directly test this, we performed phylotranscriptomic analyses of a flower development data set we previously generated (Ryan et al. 2015). Flower development follows floral transition and is dominated by the formation of different types of floral organs. In agreement with the idea that hourglass patterns in plants are not tightly associated with organogenesis, the transcriptomic profile across 14 time points from the earliest stages of flower development to mature flowers did not show an hourglass pattern or, in fact, any other pattern at all (flat line test (Drost et al. 2015a): P = 0.202, Figure 4a,b). Likewise, old and young genes did not show a clear antagonistic behavior in their expression (Fig. 4c). Together, these data suggest that in plants organogenesis is not the driving factor of hourglass-shaped transcriptome profiles. Hence, the currently favored explanation of animal hourglass patterns, which is based on selective constraints correlated to body plan establishment and organogenesis (Raff 1996), cannot serve as a plausible explanation for the two post-embryonic hourglass patterns reported here.

**Figure 4.**
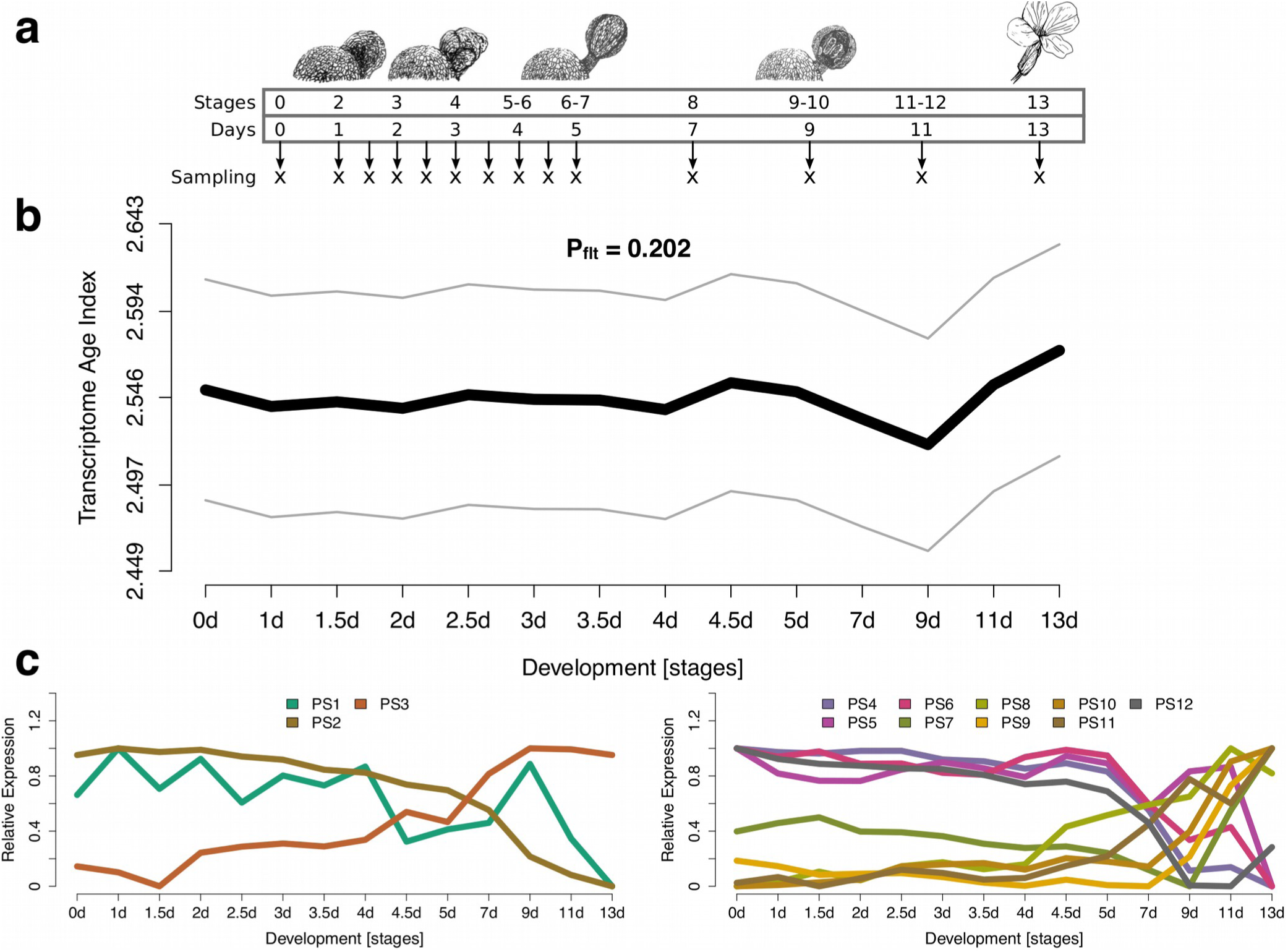
Transcriptome age index analysis of flower development in *A. thaliana*. (**A**) Illustration of the developmental stages for which transcriptome data were generated. Stages according to (Ryan et al. 2015). (**B**) The transcriptome age index (TAI) profile across flower development fails to detect evolutionary signal. The gray lines represent the standard deviation estimated by permutation analysis. The P-value was derived by application of the flat line test (Drost et al. 2015a) (P_flt_). (C) Relative expression levels for each phylostratum (PS) separately. The stage with the highest mean expression levels of the genes within a PS was set to relative expression level = 1, the stage with the lowest mean expression levels of the genes within a PS was set to relative expression level = 0, the remaining stages were adjusted accordingly. PS were classified into two groups: Group ‘old’ contains PS that categorize genes that originated before complex/multicellular plants evolved (PS1-3) and Group ‘young’ contains PS that categorize genes that originated after complex plants evolved (PS4-12).

A simple scenario that might resolve this controversy would be that the transcriptomic hourglass patterns in plants are functionally unrelated to those of animal embryogenesis. They might in fact have evolved to serve a completely different, yet unknown, purpose. This scenario is supported by the lack of reports on morphological hourglass patterns for plant embryogenesis (in contrast to various animal phyla). It seems that morphological similarity among flowering plants is not restricted to a mid-embryonic period but rather exists throughout embryogenesis (Kaplan and Cooke 1997). If the biological processes underlying embryonic hourglass patterns in animals and plants are indeed functionally unrelated, we would also have to revoke our earlier hypothesis that the developmental hourglass pattern evolved convergently in both kingdoms (Quint et al. 2012). Interestingly, in the three processes we analyzed, it seems that the waist in the hourglass reflects a general transition to a growth or maturation phase.

If, however, animal and plant hourglass patterns should serve a similar function, this study would suggest that the underlying cause is not organogenesis or body plan establishment but an even more fundamental process. As also in animal systems a causal relationship between body plan establishment and the phylotypic period remains to be proven (Irie and Kuratani 2014), it might be worthwhile to directly address this relationship by designing experiments that separate developmental transitions from organogenesis in animals.

In summary, the hourglass pattern was historically associated with animal embryogenesis and only recently recognized to govern plant embryogenesis, too. Here, we present evidence that in plants the hourglass pattern is probably even more fundamental and not only characteristic for embryo development, but present in all three major developmental transitions of plant life. It will be interesting to test post-embryogenic transitions like metamorphoses in animals to see whether this can also be observed for non-plant organisms. We hypothesize that a transcriptomic hourglass pattern is a feature of multiple developmental processes that simply require passing through an organizational checkpoint serving as a switch that separates two functional programs.

## Supplementary Material

Supplementary text, figures S1-S5, and dataset S1 are attached.

## Acknowledgements

We thank the Deutsche Forschungsgemeinschaft (Qu 141/6-1, Qu 141/7-1, and GR 3526/7-1), the Leibniz Association, the SKWP research foundation, the Science Foundation Ireland, the Netherlands Organization for Scientific Research, the Dutch Technology Foundation, which is the applied science division of the Netherlands Organization for Scientific Research and the Technology Program of the Ministry of Economic Affairs, and the “Coordenação de Aperfeiçoamento de Pessoal de Nível Superior” for financial support.

